# State-dependent relationship between lower and higher order networks in fMRI brain dynamics

**DOI:** 10.1101/2025.03.14.638301

**Authors:** Shreyas Harita, Davide Momi, Zheng Wang, John D. Griffiths

## Abstract

Resting-state functional magnetic resonance imaging (fMRI) is a powerful tool for exploring the brain’s functional organization. Functional connectivity (FC) is a commonly studied feature of fMRI data defined as the temporal correlation between activity patterns in pairs of brain regions. A major discovery of the past two decades of FC research has been the identification of consistent modular groupings in brain region time series correlations, commonly known as resting-state networks (RSNs). A second major discovery is the observation that RSNs in cortex are organized spatially along a functional/anatomical gradient. At one end of this gradient are ‘lower order’ networks (LONs), predominantly specialized for unimodal information processing. At the other end are ‘higher order’ networks (HONs), responsible for integrating multimodal information. Unlike the stable structural connectivity (SC) based on fixed anatomical links, FC fluctuates over time, and varies across brain regions. FC coordination within RSNs depends on SC, forming interconnected networks that regulate cognition, emotion, and behavior. The aim of the present study was to understand better how RSNs interact and communicate, based on their underlying SC. We used a whole-brain connectome-based neural mass modelling approach to study resting-state and task-based fMRI FC data. Following virtual SC lesions in the model, we characterized the FC changes within and between RSNs, and observed how these changes varied across different cognitive states. Our findings reveal how FC dynamics depend on underlying SC, highlighting the flexibility of these interactions across different brain states. LON lesions generally decrease FC within and between other LONs, and vice-versa for HON lesions. At rest, we observed a mutual antagonism between LONs and HONs, which was reversed during task conditions, with certain tasks increasing coordination between LONs and HONs. These results highlight the dynamic nature of brain network interactions, influenced by brain states and task demands. Our findings also have implications for clinical practice, offering insights into conditions such as brain tumors and stroke.

**Highlights:**

- The reduced Wong-Wang neural mass model was implemented in a PyTorch environment and used to understand what drives resting-state network functional connectivity (FC).
- Structural connectivity (SC) lesions were used to isolate functional networks to see how functional networks interact in various brain states.
- There is a mutual antagonism between LONs and HONs in the resting-state.
- This antagonism is seen to reverse in tasks with higher cognitive load (working memory, language).
- Tasks with a lower cognitive load (motor) share a similar network interactivity profile to the resting-state following network SC lesions.

## Introduction

### Human Brain Network Organization

Over the last two decades, resting-state functional magnetic resonance imaging (rsfMRI) has emerged as a powerful neuroimaging technique used to investigate the brain’s intrinsic functional architecture. By measuring spontaneous fluctuations in blood oxygen level-dependent (BOLD) signals, rsfMRI provides insights into functional connections between different brain regions, even without task-based inputs or external stimuli (Lee et al., 2013). The primary method for investigating intrinsic neural fluctuations involves assessing correlations among voxel- or region-specific time series data obtained through rs-fMRI, commonly known as “functional connectivity” (FC). The initial observation of strong rs-fMRI FC between hemispheric homologues in the primary motor cortex by Biswal et al. in 1995 marked the inception of this field (Biswal et al., 1995). Subsequently, extensive research within the neuroimaging community has unveiled the presence of distinct ‘functional networks’ distributed throughout the brain. These networks, often called resting-state networks (RSNs), represent prominent shared variance components within rs-fMRI data, highlighting the intricate connectivity patterns underlying brain function. RSNs are characterized by their robust patterns of FC with and between multiple anatomically distinct regions (M. D. Fox & Raichle, 2007). Research has identified several distinct RSN groups, with the most prominent including the Visual Network (VN), Somatomotor Network (SMN), Dorsal Attention Network (DAN), Ventral Attention Network (VAN), Limbic Network (LN), Frontoparietal Network (FPN), and Default Mode Network (DMN) (Schaefer et al., 2018; Yeo et al., 2011). These networks demonstrate consistent characteristics across diverse populations and under different fMRI protocols (Rosazza & Minati, 2011). Of particular significance is the observation that many of the brain regions underlying RSNs also correspond to established functional networks associated with specific tasks (van den Heuvel & Hulshoff Pol, 2010). An important feature of RSNs discovered in recent years (Margulies et al., 2016) is that they fall in a consistent spatial arrangement along the first non-trivial eigenvector of rsfMRI FC, also known as the ‘principal gradient’. On one end of this gradient lie networks implicated in unimodal information processing, including the VN, SMN, DAN, and VAN. The other end of the gradient contains networks that process multimodal information, including the DMN, FPN, and LN (Margulies et al., 2016). The unimodal networks are concerned with perceiving information and responding to it. In the present paper we refer to those networks that largely process and respond to external stimuli/sensory information as ‘lower order’ networks (LONs), and those mainly concerned with integrating multiple information processing streams as ‘higher order’ networks (HONs).

### The Relationship Between Functional and Structural Connectivity

Structural connectivity (SC), defined by the presence and strength of anatomical connections, necessarily remains stable over short time intervals. In contrast, FC exhibits substantial variation across various spatial and temporal scales. This variability contributes to the brain’s dynamic and diverse activity patterns. Importantly, the coordination of neural activity, which FC is believed to be reflective of, has to be mediated, ultimately, by transmission of signals through SC. Research linking brain structure and function is driven by the understanding that the human brain’s activity relies on the network of neurons and their connections. Thus, SC and FC are inherently linked. Evidence supporting this has shown systematic coupling of lifespan changes in SC and FC (Baum et al., 2020; Romero-Garcia et al., 2014). Additionally, studies have found a notable similarity between white matter connectivity profiles and functionally significant cortical parcellations (Greicius et al., 2009; Jung et al., 2017; Vázquez-Rodríguez et al., 2019). Research has also shown that highly central nodes within functional networks have strong, direct white matter connections between them (Greicius et al., 2009). Evidence suggests that the brain’s SC-FC relationship is shaped by the spatial organization of neuronal populations, which determines activation sites and is influenced by cortical properties such as surface area, volume, curvature, and thickness (Litwińczuk et al., 2022). Local connectivity density affects the amount of cortical activity exchanged nearby (Bassett & Bullmore, 2017; Bullmore & Sporns, 2012), while efficient activity exchange across distant regions is facilitated by the integrity of long-range white matter tracts (Bullmore & Sporns, 2012; Liu et al., 2017). Based on the above evidence, although certain fundamental aspects of the SC-FC relationship are becoming clearer, our precise comprehension of the complex interplay between brain structure and function remains incomplete.

### Lesioning brain structure to understand brain function

As we explore the intricate relationship between SC and FC, we acknowledge that disruptions to these networks can yield valuable insights into brain function. Brain lesions offer a unique opportunity to probe the functional consequences of localized damage to specific brain regions, resulting from various causes such as injury, stroke, infection, or neurodegenerative diseases (Nabizadeh & Aarabi, 2023). Historically, the study of brain lesions and their associated deficits, known as lesion mapping, has provided invaluable insights into the localization and functional specialization of different brain regions. Lesion studies have been instrumental in identifying brain regions responsible for language processing, motor control, sensory perception, memory, and various other cognitive functions (Rorden & Karnath, 2004).

From a network neuroscience perspective, and with the advent of modern neuroimaging modalities, lesion mapping has evolved to correspond to the weakening or removing of network nodes and/or edges from the structural connectome. Modern lesion studies are carried out in a model of the human (or animal) connectome. This ‘network lesioning’ has been and continues to be a valuable way of studying neuropathologies that involve real lesions. However, for the purposes of the present work, it is important to note that network lesioning can also be used as a theoretical tool to understand aspects of network organization, including brain network organization, independently of any specific reference to real brain lesions per se. Examples from previous research have looked at the effects of lesioning specific nodes within a functional network and its impact on, both, FC within the network the node belonged to, as well as on the FC of distant nodes away from the lesion. In general, these studies report that there is a local (i.e., within the network) and global (i.e., away from the lesion) reduction in FC when nodes with high degree of centrality are removed (Alstott et al., 2009; Honey & Sporns, 2008; Váša et al., 2015).

### Present Study

In the present study, we sought to understand how the various functional networks of the brain communicate based on the underlying SC. Our methodological approach involved using whole-brain connectome-based neural mass modeling of rs-fMRI FC data. Neural mass models consider the average activity of an entire neuronal population, representing a brain area or parcel, rather than individual neurons within that brain region. Connectome-based brain network models integrate the activity of multiple neural masses based on SC, typically derived from diffusion-weighted MRI (DW-MRI) tractography (Breakspear, 2017). This approach has been utilized to explore resting-state brain activity (Deco et al., 2011, 2013; Honey et al., 2009), brain oscillations (Breakspear et al., 2010), alpha waves (Griffiths et al., 2020), and traveling waves (Breakspear, 2017; Heitmann et al., 2012), among other phenomena. Here, we constructed a connectome-based brain network model utilizing the reduced Wong-Wang (RWW) neural mass model, and implemented it in the Whole Brain modelling in PyTorch (WhoBPyT) software library (Griffiths et al., 2022; Momi et al., 2023), to study network interactions and how they change following SC lesions in resting-state fMRI data. The RWW model has been widely employed in previous research to investigate both resting-state and task-evoked brain dynamics, with a particular focus on rs-fMRI activity, for which it was specifically designed (Deco et al., 2013, 2014).

To investigate the interactions between functional networks in relation to SC, we employed the latter use of the lesion methodology discussed above. Virtual lesions were created to isolate different functional networks structurally, allowing us to observe changes in FC within and between these networks. Our hypothesis posited that lesions to network SC would reduce FC both within and between the lesioned and other functional networks, with the extent of FC changes depending on the brain’s current state. To validate this hypothesis, we examined the effects of network SC lesions not only during resting-state conditions but also in task-based fMRI data across motor, working memory (WM), and language tasks. A comprehensive understanding of functional network interactions across different brain states holds promise for clinical applications and sheds light on the complexities of the SC-FC relationship.

## Methods

The methodology used in this study includes the following components: a) determining the SC matrix using diffusion-weighted MRI tractography data, b) simulating whole-brain resting-state and task fMRI time series with a neurophysiological model based on the SC matrix, c) optimizing model parameters using a PyTorch-based fitting approach to match the simulated and subject-specific fMRI time series from the Human Connectome Project (HCP) database, and d) systematically creating virtual lesions in canonical functional networks to simulate ‘lesioned’ fMRI time series and assess the impact of SC lesions on functional connectivity within and between networks (Figure 1).

**Figure 1:**
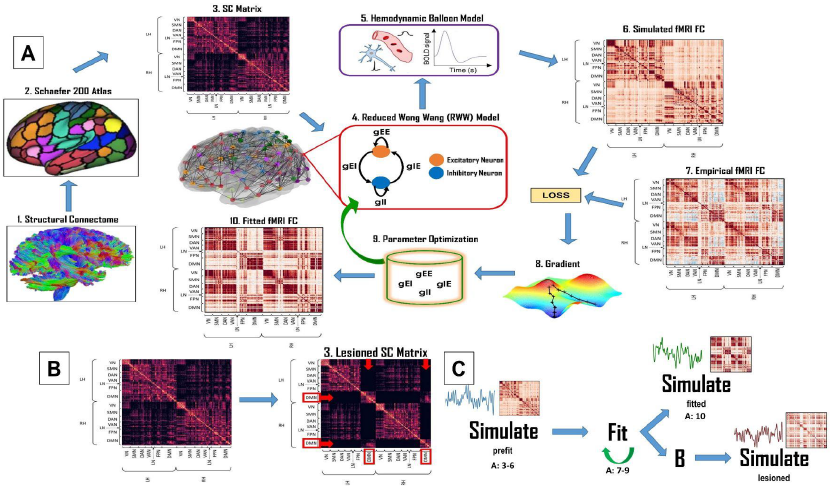
Overview of the methodological approach. **A - Simulating fMRI data. 1-2:** We use DW-MRI tractography data to determine the SC matrix for the whole-brain model which contains the RWW model equations at each node. The nodes are based on the brain parcellations determined from the Schaefer atlas (200 regions). **3-6:** The RWW model, together with the Balloon-Windkessel model, are then used to simulate resting-state and task-based fMRI FC. **7-9:** We optimize the parameters of our model using a gradient-based parameter optimization scheme utilizing automatic differentiation-based algorithms. **10:** This allows us to maximize the fit between simulated and empirical fMRI FC data, resulting in the fitted fMRI FC matrix. **B - Creating virtual SC lesions**. Having obtained the simulated fMRI FC from the intact SC, we then virtually lesioned the SC to structurally isolate a given network. In this example, we see the DMN has its connections removed (red arrows) to all the other networks, except its own cross-hemispheric homologues. **C - Flowchart of methodological approach**. We begin by simulating the FC time series/matrix prior to fitting any model parameters. We then iteratively optimize our model parameters (‘fitting’) to obtain the fitted FC time series/matrix. Following the virtual SC lesion, we again simulate using the previously fitted parameters to obtain the lesioned FC times series/matrix.

### Structural Connectivity Data

DW-MRI data were obtained from a cohort of 200 (120 female) randomly selected subjects in the HCP WU-Minn cohort database (Van Essen et al., 2013). The preprocessing pipeline was executed on Ubuntu 18.04 LTS and utilized tools from the FMRIB (FSL 5.0.3; www.fmrib.ox.ac.uk/fsl; (Jenkinson et al., 2012)), MRtrix3 (www.MRtrix.readthedocs.io) (J.-D. Tournier et al., 2012) and FreeSurfer (Fischl, 2012) software libraries. Motion correction was performed using FSL’s EDDY (Andersson & Sotiropoulos, 2016) as part of the HCP minimal preprocessing pipeline (Glasser et al., 2013). The multi-shell multi-tissue response function (Christiaens et al., 2015) was estimated using a constrained spherical deconvolution algorithm (Jeurissen et al., 2014). The T1w images, already coregistered to the b0 volume, were segmented using the FAST algorithm (Zhang et al., 2001). Anatomically constrained tractography generated the initial tractogram with 10 million streamlines (Amico et al., 2017) using second-order integration over fiber orientation distributions (J. Tournier et al., 2009). This was followed by the application of the Spherical-deconvolution Informed Filtering of Tractograms (SIFT2) methodology (Smith et al., 2015) for more biologically accurate measures of fiber connectivity.

Following tractography reconstructions, the gray/white matter interface underwent anatomical parcellation to segment the whole-brain streamline sets into pairwise bundles, yielding a SC matrix for each subject. The parcellation used was the 200-node variant of the brain atlas (Schaefer et al., 2018), which assigns each parcel to one of the 7 canonical functional networks identified by Yeo et al., (2011) - VN, SMN, DAN, VAN, LN, FPN, and DMN. Parcels were defined at the surface vertex level, aligned with each individual’s FreeSurfer reconstruction via spherical registration (Fischl, 2012), and transformed into native-space image volumes registered with the DW-MRI tractography streamlines. Entries in the resulting 200x200 SC matrices for each subject denote the number of white matter streamlines connecting each pair of ROIs.

### Resting-State and Task Empirical fMRI data

The resting-state and task-based fMRI data used in this study was obtained from 200 (120 female) randomly selected subjects from the HCP database. These conditions are briefly described below. In the resting-state, the participant’s eyes were open and they were instructed to maintain a relaxed fixation on a white cross on a dark background, thinking about nothing specific, without falling asleep. Additionally, task-based fMRI data for the same 200 subjects were obtained for the motor, WM, and language tasks. For the motor task, visual cues guided participants to either tap their left or right fingers, squeeze their left or right toes, or move their tongue. For the WM task, participants performed a visual n-back task, with 0- and 2-back conditions, along with four stimulus categories (place, tools, faces, body parts) which were also blocked. For the language task, the participants engaged in aurally presented alternating story and math tasks. For the complete HCP acquisition protocols, and other related information, please refer to Glasser et al., (2013), Uğurbil et al., (2013), Van Essen et al., (2013), and Van Essen et al., (2012). For more detailed task descriptions, see Barch et al., (2013).

### Neurophysiological Model

#### Dynamic Mean Field Model

To capture mesoscopic population activity across various brain regions within a whole-brain network, we employ a mathematical model known in the literature as the ‘Dynamic Mean Field’ model, or ‘Reduced Wong-Wang’ (RWW) model (Deco et al., 2013, 2014; Demirtaş et al., 2019; Hansen et al., 2015; Pang et al., 2022). Specifically, we use the two-state or E-I extension (Deco et al., 2014) of the RWW equations first introduced by Deco and colleagues (Deco et al., 2013), based on the original mean-field model of Wang and colleagues (Wong & Wang, 2006). This model characterizes each node in the brain network with two neural masses, one representing the behavior of an excitatory neural subpopulation and the other describing the inhibitory subpopulation. The activities of these subpopulations at a given brain network node i ∈ [1…N] are described by unitless state variables 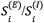,which represent synaptic activity or ‘gating’ levels. Additionally, auxiliary variables 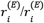 denote population firing rates. These variables evolve according to a set of coupled nonlinear stochastic differential equations:

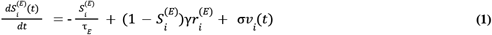

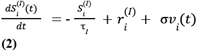

Here, τ*E* and τ1 are the decay times for excitatory and inhibitory synapses, γ is a kinetic parameter (see *Supplementary Information - Table 1* for values), and σ*ν*_*i*_ is uncorrelated Gaussian noise with a mean of 0 and standard deviation of σ. The principal inputs to 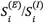 are population firing rates 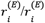, expressed as functions *H* of the synaptic input currents 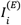 and 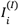. These relationships are defined as follows:

**Table 1:** Default values for the RWW model. These are the values used for the constant parameters of the RWW model and are identical to the ones described in Deco et al., (2014).

| Parameter Name | Description | Symbol | Value |
| --- | --- | --- | --- |
| Excitatory parameter values | The excitatory parameter values for the input-output function helps convert incoming inputs to the excitatory population into firing rates. | $\mathbf{a_E}$ | 310 $\text{nC}^{-1}$ |
| | | $\mathbf{b_E}$ | 125 (Hz) |
| | | $\mathbf{d_E}$ | 0.16 (s) |
| Inhibitory parameter values | The inhibitory parameter values for the input-output function helps convert incoming inputs to the inhibitory population into firing rates. | $\mathbf{a_I}$ | 615 $\text{nC}^{-1}$ |
| | | $\mathbf{b_I}$ | 177 (Hz) |
| | | $\mathbf{d_I}$ | 0.087 (s) |
| Excitatory time constant | Decay time constant for NMDA receptors. | $\tau_E$ | 100 ms |
| Inhibitory time constant | Decay time constant for GABA receptors. | $\tau_I$ | 10 ms |
| Excitatory scaling weight | Scales the external input to the excitatory pool. | $\mathbf{W_E}$ | 1 |
| Inhibitory scaling weight | Scales the external input to the inhibitory pool. | $\mathbf{W_I}$ | 0.7 |
| Kinetic parameter | Expressed in milliseconds by dividing by 1000. | $\gamma$ | 0.641/1000 |

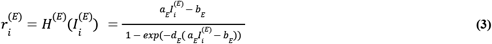

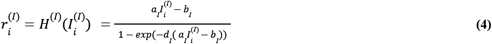

The parameters *a*_*E*_, *b*_*E*_, *d*_*E*_, *a*_*I*_, *b*_*I*_, *d*_*I*_ are constant values that help to convert input currents into population firing rates (see *Supplementary Information - Table 1* for values). The synaptic input currents are computed as follows:

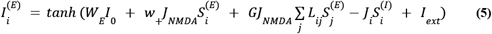

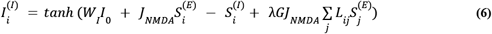

Here, *I*_*ext*_ encodes external stimulation and is set to 0 for simulating resting state activity. *I*_0_ represents a constant external input (set to 0.2 nA), scaled by parameters *W*_*E*_ and *W*_*I*_ for the excitatory and inhibitory populations, respectively. *L*_*ij*_ represents the elements of the connectivity Laplacian, calculated as *L*_*ij*_ = *D* − *C* where *C* is the tractography-derived (see above) structural connectivity matrix that denotes the connection strength between network nodes *i* and *j*, and *D* is the diagonal matrix of the row sums of C (node degree). The term 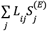 encodes the total summed input to node *i* from all other *j* nodes in the network. The parameter λ, when set to 0, allows the removal of long-range feedforward inhibition (Deco et al., 2014). The *J*_*NMDA*_ and *J*_*i*_ parameters represent the values of the excitatory and inhibitory synaptic coupling, while parameters *w*_+_ and *G* scale the local and long-range excitatory couplings, respectively.

An important feature of the RWW model is its derivation from a lower-level mathematical description of individual neurons, specifically conductance-based leaky integrate-and-fire neurons. The resultant two-state RWW model closely resembles the classic equations of Wilson and Cowan (1972), which describe excitatory/inhibitory population interactions in cortical tissue as a predator/prey-like system. In addition to this, Deco et al., (2014) introduced an algorithm to maintain synaptic current within a biologically motivated range, and in this context, we constrained the input current variables in Eqns. 5-6 using a *tanh* function. This choice also has the important additional benefit of providing better model fit performance compared to explicit constraints or iterative algorithms.

While the majority of the above notations are identical to those in Deco et al. (2014), here we adopt an alternative notation for key terms as follows:

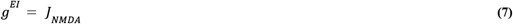

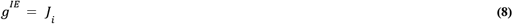

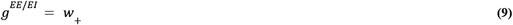

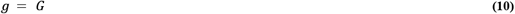

These notations define the within-node excitatory-to-inhibitory (*g*^*EI*^), inhibitory-to-excitatory (*g*^*IE*^), excitatory-to-excitatory (*g*^*EE*^) synaptic gains, and the long-range global coupling (*g*) (see *Supplementary Information - Table 2* for value ranges). These four parameters will be altered by the neural network (see *Optimization of Dynamic Mean Field Model Parameters for Individual Subjects* below) by comparing model simulations to empirical data to achieve an optimal fit.

**Table 2:** Estimated parameter values for the RWW model for resting state and task simulations. This table summarizes the prior and posterior values of the fitted variables used in the RWW model. Each column under the posterior values tab shows a column for each condition. Within each condition’s column the mean ± standard deviation is highlighted, along with the range of values in italics below. This table shows the average prior and posterior values across all 200 subjects. Notable features of the estimated parameters are as follows: 1) Three of the four (g^EE^, g^EI^, g^IE^) parameters change moderately from their prior values. However, the value they move to is largely consistent across all four conditions. 2) The largest difference across conditions is between rest and the three tasks. This is particularly noticeable for the g^EE^ and g^EI^ parameters.

| Fitted Variables | Prior Values | Posterior Values |  |  |  |
| --- | --- | --- | --- | --- | --- |
|  |  | Resting-State | Motor Task | WM Task | Language Task |
| $g$ | 400 | **$400.01 \pm 0.16$** | **$399.99 \pm 0.16$** | **$400.01 \pm 0.17$** | **$400.00 \pm 0.16$** |
|  |  | *Range: 399.53 - 400.46* | *Range: 399.55 - 400.48* | *Range: 399.57 - 400.39* | *Range: 399.64 - 400.44* |
| $g^{EE}$ | 1.5 | **$1.59 \pm 0.08$** | **$1.51 \pm 0.07$** | **$1.53 \pm 0.07$** | **$1.52 \pm 0.07$** |
|  |  | *Range: 1.36 - 1.73* | *Range: 1.27 - 1.74* | *Range: 1.30 - 1.67* | *Range: 1.30 - 1.68* |
| $g^{EI}$ | 0.8 | **$0.99 \pm 0.41$** | **$0.73 \pm 0.19$** | **$0.75 \pm 0.22$** | **$0.71 \pm 0.21$** |
|  |  | *Range: 0.24 - 1.99* | *Range: 0.24 - 1.19* | *Range: 0.18 - 1.38* | *Range: 0.20 - 1.22* |
| $g^{IE}$ | 0.6 | **$0.47 \pm 0.03$** | **$0.48 \pm 0.02$** | **$0.48 \pm 0.02$** | **$0.49 \pm 0.02$** |
|  |  | *Range: 0.41 - 0.53* | *Range: 0.44 - 0.54* | *Range: 0.44 - 0.54* | *Range: 0.44 - 0.56* |

#### Balloon Windkessel Model

The balloon-windkessel (or ‘balloon’) hemodynamic model is used to relate changes in the BOLD signal to changes in blood flow and oxygenation. The ‘balloons’ in the model are the blood vessels in the brain, which can expand and contract with changes in blood flow. The model also takes into account the rate at which oxygen is consumed by the active brain tissue, and the rate at which oxygen is delivered by the blood. The following equations briefly describe the balloon model and how it relates to the neuronal signal.

The signal that induces blood flow is generated by neuronal activity *u*(*t*), where *u*(*t*) is 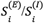 (*Eqs. 1 and 2*) for excitatory and inhibitory neuronal activity, respectively.

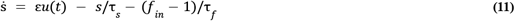

where ε, τ_*s*_, and τ_*f*_ have typical values associated with them (*See Supplementary Information - Table 3 for values*).

**Table 3:** Default values for the Balloon model. These are the values used for the constant parameters of the Balloon model. These values are similar to the ones described in Friston et al., (2000, 2003).

| Parameter | Description | Symbol | Value |
| --- | --- | --- | --- |
| Signal decay | Describes the rate at which the BOLD signal is eliminated, helping to accurately model the hemodynamic response. | $\tau_s$ | 1.54 |
| Transit time | Describes the time it takes for blood to travel through the venous compartment. | $\tau_0$ | 0.98 |
| Autoregulation | This parameter represents the time constant of the feedback autoregulatory mechanism, though its physiological nature remains unspecified. | $\tau_f$ | 2.44 |
| Stiffness | Represents the elasticity or compliance of the venous compartment. | $\alpha$ | 0.32 |
| Neuronal efficacy | Represents the increase in perfusion signal elicited by neuronal activity. | $\epsilon$ | 0.5 |
| Resting oxygen extraction | Refers to the proportion of oxygen extracted from the blood by tissues under resting conditions. | $E_0$ | 0.34 |
| Resting blood volume fraction | Refers to the fraction of blood volume in the brain's vascular system. | $V_0$ | 0.02 |

The inflow of blood *f*_*in*_ is also related to the signal generated by *u*(*t*):

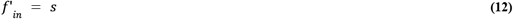

With the inflow of blood, there is the extraction of oxygen from the hemoglobin and the outflow of blood *f*_*out*_ with the change in deoxyhemoglobin (*q*/*q*’).

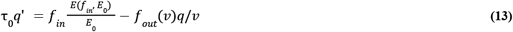

where, *E*(*f*_*in*_, *E*_0_) is the fraction of the oxygen extracted from the inflowing blood, τ_0_ is the time constant, and *v* is the volume of venous blood. The rate of change of *v* is given by

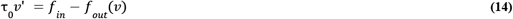

In Eqs. 13 and 14, *q* represents the normalized total amount of deoxyhemoglobin within the voxels (with q = 1 at rest), and *v* denotes the normalized volume of venous blood (with v = 1 at rest) (Buxton et al., 1998).

The BOLD signal *y*(*t*) is a static nonlinear function of normalized *v*, normalized total *q*, and the resting net oxygen extraction fraction (*E*_0_), given by the equation

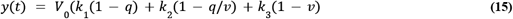

where *k*_1_ = 7*E*_0_, *k*_2_ = 2, *k*_3_ = 2*E*_0_ − 0. 2, and *V*_0_ is the resting blood volume fraction.

#### Optimization of Dynamic Mean Field Model Parameters for Individual Subjects

For this study, we employed a recently described brain network model parameter optimization technique as detailed in Griffiths et al., (2022) and Momi et al., (2023), and extended in Clappison (2024). A notable aspect of this approach is the utilization of PyTorch (Paszke et al., 2019), a software library that has gained widespread acceptance in both academic and commercial sectors within the machine learning community in recent years. Transitioning to PyTorch from more traditional numerical simulation libraries required some minor adjustments to accommodate tensor data structures. However, this shift offered a significant advantage by naturally supporting gradient-based parameter optimization through automatic differentiation-based algorithms. This was particularly valuable for handling complicated sets of equations that do not readily yield quantifiable Jacobians (Griffiths et al., 2022). This application underscores the increasing convergence between our physiologically-based large-scale brain network models and the deep recurrent neural networks employed in machine learning, proving to be both practically and conceptually beneficial, as demonstrated in related works such as Richards et al., (2019).

The full brain network model, which consists of the dynamic mean field model (*equations 1-10*) and the balloon windkessel model (*equations 11-15*), can be likened to a form of recurrent artificial neural network (ANN). It utilizes parameters like *C*_*ij*_ (which, in turn, define *L*_*ij*_), *g*^*EI*^, *g*^*IE*^, *g*^*EE*^, and *g* as its weights. Among these, the *C*_*ij*_ parameters are especially comparable to the free parameters in an ANN, as they define the strengths of connections between individual network nodes. In contrast, the *g* parameters have a broader impact since they scale the connection strengths among all nodes in the network simultaneously. All other physiological and hemodynamic model parameters are considered fixed and are not discussed here.

To estimate these model parameters, we employ a machine learning-based optimization approach. We achieve this by aligning the activity generated by the model with empirical functional neuroimaging data. To dynamically fit the brain network model to the empirical neuroimaging data, we divide the fMRI BOLD time series into non-overlapping 30-second windows (batches). Within each batch, we generate a simulated BOLD time series using our model, taking into account the current estimated parameter values. We calculate a cost to guide updates of the model parameters for the subsequent batch. Our goal is to discover optimal values for parameters *C*_*ij*_, *g*^*EI*^, *g*^*IE*^, *g*^*EE*^, and *g* in Equations 7-10 to minimize the objective function.

For the parameter optimization process, we employ the ADAM algorithm (Kingma & Ba, 2014), natively supported in PyTorch. ADAM is an adaptive learning rate optimization algorithm specifically designed for training deep neural networks. It combines elements of RMSprop (Tieleman & Hinton, 2012) and Stochastic Gradient Descent (SGD) with momentum. ADAM adjusts the learning rate using squared gradients (similar to RMSprop) and leverages momentum by using the moving average of gradients instead of the gradients themselves (similar to SGD with momentum). This adaptability is achieved by estimating the first and second moments of the gradient of the model parameters concerning the objective function and using this information to adapt the overall learning rate. The general mathematical framework underlying this approach is elaborated in a recent technical paper by Griffiths et al., 2022. In the present study, this approach was applied in the context of connectome-based neurophysiological modeling using resting-state and task-based fMRI data.

### Structural connectivity lesions to isolate functional networks

Following parameter optimization using the ADAM algorithm, and ensuring that our model was able to sufficiently replicate the rs-fMRI times series, we conducted a series of numerical simulation experiments where the SC is virtually lesioned, so as to isolate the seven canonical functional networks described above. This approach provides a means of studying how functional networks (i.e., the FC patterns within the system) behave when structurally isolated from each other, and how this affects the relationship between the other non-lesioned functional networks. We applied this network isolation methodology to simulations of both resting-state and task-based fMRI activity, giving us insight into the differences in functional network organization across different neurocognitive states. We studied the FC within a network (i.e., the FC of a network to itself) and the FC between a given network and other networks, following the SC lesion of the target network. This process was repeated for each of the seven networks for each condition. We describe the results of SC lesions based on their effects on LONs, HONs, and between LONs and HONs. These effects are reported as percentage changes in the average FC of each network to the others when one of the seven networks is lesioned. The percentage changes are depicted as 7×7 symmetrical matrices. The elements of each grid show the percentage change in the FC between the seven functional networks, with the diagonal of each matrix representing the percentage FC change within each network.

### Statistical Analysis

Pairwise t-tests were conducted between the intact and lesioned SC average FC. Our alpha threshold (p-value) was set to be less than 0.01 and only significant percent changes in FC are reported in the results. Additionally, we conducted a permutation analysis to examine the significance of percent change patterns observed in network FC within and between functional networks. Permutation analysis is a statistical technique used to determine the significance of observed patterns in data by comparing them against a distribution of patterns generated under a null hypothesis. This method involves randomizing or ‘shuffling’ the data to create a large number of permutations, thereby establishing a distribution of outcomes that would be expected by chance. The results of this analysis can be found in the *Supplementary Information* section.

## Code and Data Availability

All analyses presented in this paper were conducted on CentOS Linux compute servers running Python 3.7.3, utilizing the standard scientific computing stack and various open-source neuroimaging software tools, primarily including, Nibabel for neuroimaging data input/output (Brett et al., 2020), and Nilearn for neuroimaging data visualization (Abraham et al., 2014). The code and analysis results are openly accessible at github.com/griffithslab/HaritaEtAl2024_whobpyt-network-sc-lesions.

## Results

Using the whole-brain computational modelling framework detailed above, we studied how SC lesions affect the simulated FC connectivity, the relations within and between the 7 canonical functional networks mentioned above in the resting state, and task-based FC paradigms. In the following, we describe the results of our simulations using the RWW model. We highlight the model fit to empirical resting-state and task-based fMRI data and use the SC lesions to understand how these networks interact in resting-state or task-based conditions.

### Model simulations reproduce primary characteristics of empirical resting-state and task fMRI FC data

For every individual within our cohort of 200 subjects, simulated resting-state BOLD FC using WhoBPyT was generated, followed by assessing the fit through Pearson correlation between the simulated and empirical FC matrix upper triangles. This process returned an average fit of R = 0.64 (Std. Dev = 0.05), ranging from R = 0.48 to R = 0.77. The simulated FC from our model can accurately capture features of the empirical FC matrix. For example, the canonical resting-state networks are replicated in the simulated data and closely align with those in the empirical data. This is seen in the left and right hemispheres (Figure 2A).

**Figure 2:**
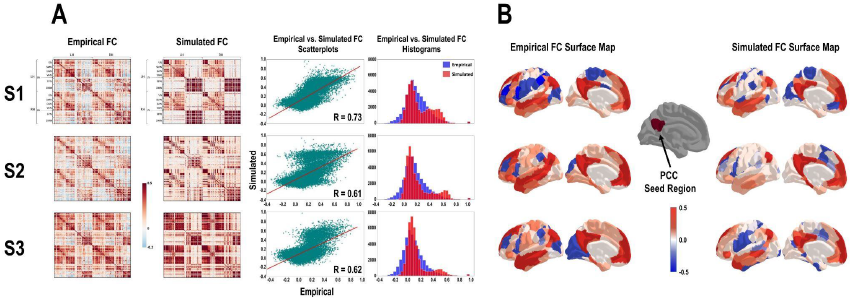
Model fit to empirical resting-state data. **A -** Goodness of fit of the RWW/WhoBPyT model. Across our subject cohort, the model showed an average fit of R=0.64 ± 0.05 to the empirical data. From left to right: heatmaps compare the empirical and simulated FC matrices; scatterplots show the fit of the simulated FC to the empirical FC; histograms show the distribution of the FC for empirical (blue) and simulated (red). **B** - Cortical surface FC maps for the posterior cingulate cortex (PCC) seed region. The patterns of FC for the PCC seed region in the *left* hemisphere are presented for the empirical (left) and simulated (right) data. Both A and B show data from same three randomly selected subjects from our 200-subject cohort.

Additionally, when examining the cortical surface FC maps, we used the posterior cingulate cortex (PCC) as a seed region to study the features of the FC simulated by our model and compare that with the empirical FC. We noticed that the model sufficiently replicates the positive correlations (PCs) in the simulations in regions t like the temporal lobe, parietal lobe, and medial prefrontal cortex. This is evident across all three randomly selected subjects in Figure 2B. However, while the PCs in the simulated FC match the empirical FC, the same cannot be said about the negative correlations (NCs). Typically, we noticed that across regions/subjects, there is some alignment of NCs with the empirical data, however, most empirical NCs are not replicated or have their FC values closer to zero in the simulated FC.

For all 200 subjects, the above process was repeated with task-based fMRI data for the motor, WM, and language tasks from the HCP database. Overall, the model fit to the task empirical task data was lower than the fit to the empirical resting-state data. The motor task had the highest average fit of the simulated to empirical data, with R = 0.53 (Std. Dev = 0.08; min. R = 0.26; max. R = 0.70). The WM task had an average fit of R = 0.49 (Std. Dev = 0.09; min. R = 0.22; max. R = 0.69). The language task had the lowest average fit of simulated to empirical data, with R = 0.31 (Std. Dev = 0.10; min. R = 0.10; max. R = 0.54).

In the FC for the task simulations, a unique seed region was chosen for each task to study how the model can capture features of task-based fMRI. For the motor task, a seed region within M1 was selected. Here we note that the model can replicate the bulk of the strong PCs seen in the lateral view of the left and right hemispheres. However, the NCs at the temporal lobes and on the left and right medial surfaces are not seen in the simulated data (Figure 3A). The dlPFC was chosen as the seed region for the WM task. Here, we noticed a good replication of the PCs seen on the lateral and medial surfaces of the left and right hemispheres. We also note the partial replication of NCs in simulated data at the right temporal pole. However, other NCs seen in the empirical data were not seen in the simulated data (Figure 3B). Lastly, for the language task, the seed region was a portion of the temporal cortex. While there are some PCs that are comparable between the empirical and simulated FCs, this task also has the lowest simulated fit to the empirical data. Thus, no other specific similarities are seen in the simulated data (Figure 3C).

**Figure 3:**
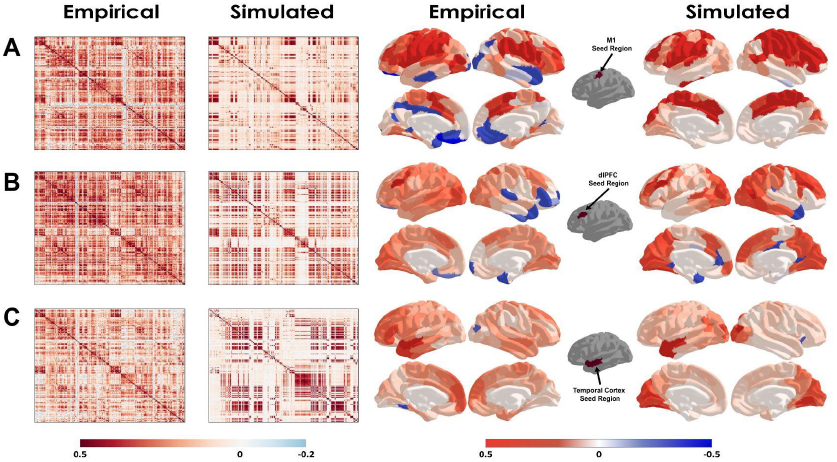
Similarities between simulated and empirical task data. Empirical and simulated FC data for the same randomly selected subject (S1 from Figure 2) for three different tasks - motor, WM, and language. Left: Heatamps of correlations for all Schafer 200 region pairs. Right: Cortical surface FC maps with a task-specific seed region for the same subject. The simulated FC data is able to captures most of the relevant features of the empirical FC data and how these features differ across various tasks.

### LON SC lesions decrease the FC in other LONs and increase the FC in HONs

Generally, following the lesion of any LON, there is a decrease in the FC within and between LONs (15.1% average reduction), with the lesioned LON typically suffering the highest FC reduction (in some cases, well over 40%). There is also a decrease in the FC between LONs and HONs (10.3% average reduction). Finally, following any LON lesion, an increase in FC is observed within and between HONs (2.5% average increase). These changes in FC are seen in the resting state (Fig. 4B [left]), motor (Fig. 5B [left]), WM (Fig. 6B [left]), and language task (Fig. 7B [left]) conditions. However, this is a generalization, and there are unique changes associated with the FC for each lesioned network.

**Figure 4:**
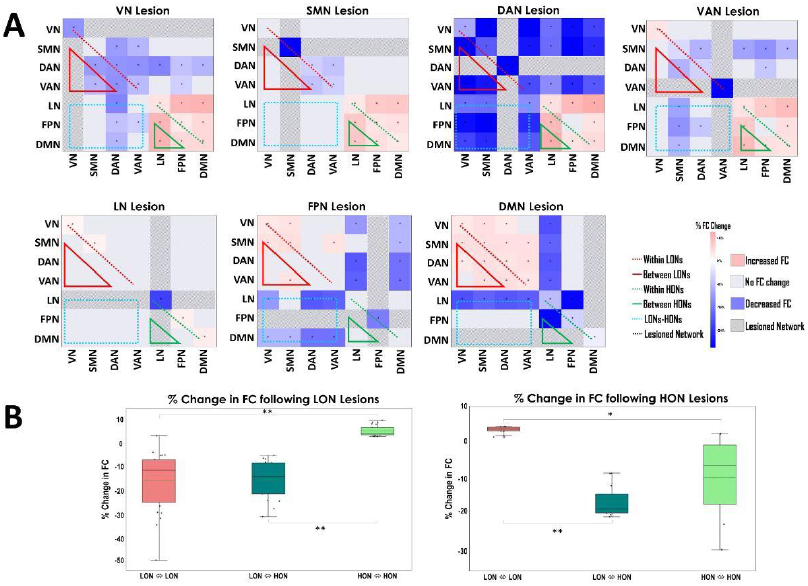
The effect of SC lesions on the FC of canonical networks in the Resting State. **A** - Each element within a 7×7 grid represents the observed changes in the average FC of the given network to the other networks (represented as % changes) when one of the seven networks is lesioned. All changes (i.e., increase or decrease) in FC reported here are significant (p < 0.01). ***Top (from left to right)***: LON lesions - VN lesion, SMN lesion, DAN lesion, VAN lesion. ***Bottom (from left to right)***: HON lesions - LN lesion, FPN lesion, DMN lesion. Mutual antagonism between LONs and HONs is evident in the consistent blocks of blue and red in the first and second rows. **B** - Box plots summarizing the overall percent change in FC following a LON lesion *(left)* and HON lesion *(right)*. For FC changes within and between LONs, the red box plot represents the percent change in FC for the networks marked with the red dotted and solid lines in panel A. For changes in FC between LONs and HONs, the teal box plot shows the percent change in FC for the networks indicated by the blue dotted line in panel A. Changes within and between HONs are represented by the pale green box plot which is the percent change in FC for the networks highlighted with the green dotted and solid lines in panel A. ‘*’ or ‘**’ is used to indicate a significant difference between the boxplots in panel B. **\*\*** = p<0.01; **\*** = p<0.05. The mutual antagonism represented in A with the patterns of red vs. blue is re-represented here primarily in the positive values of the 3rd and 1st bars of the left and right box plots respectively.

**Figure 5:**
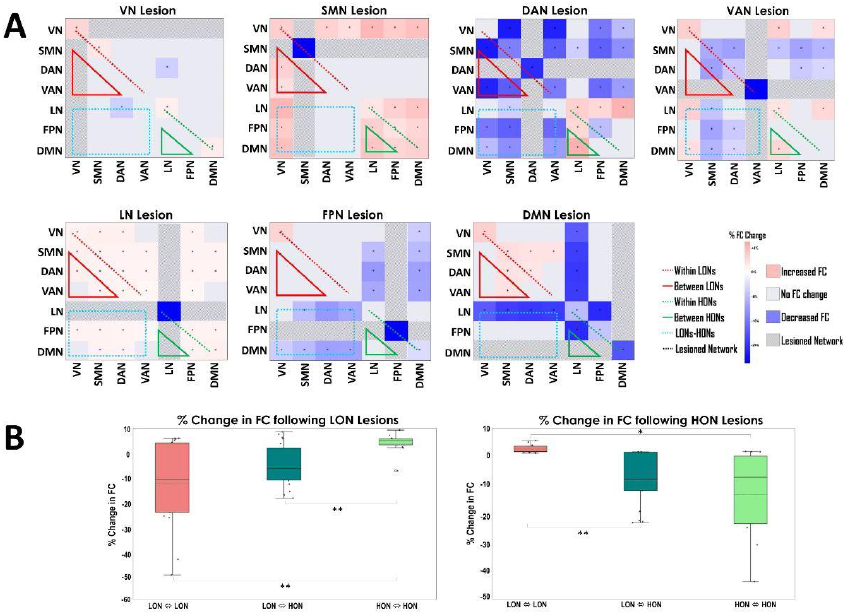
The effect of SC lesions on the FC of canonical networks for the Motor Task. The legend for this figure is identical to Figure 4 for parts A and B. In this figure we can see the patterns of connections between LONs and HONs being similar to the resting-state paradigm above. This is potentially due the lower cognitive load imposed by the motor task.

**Figure 6:**
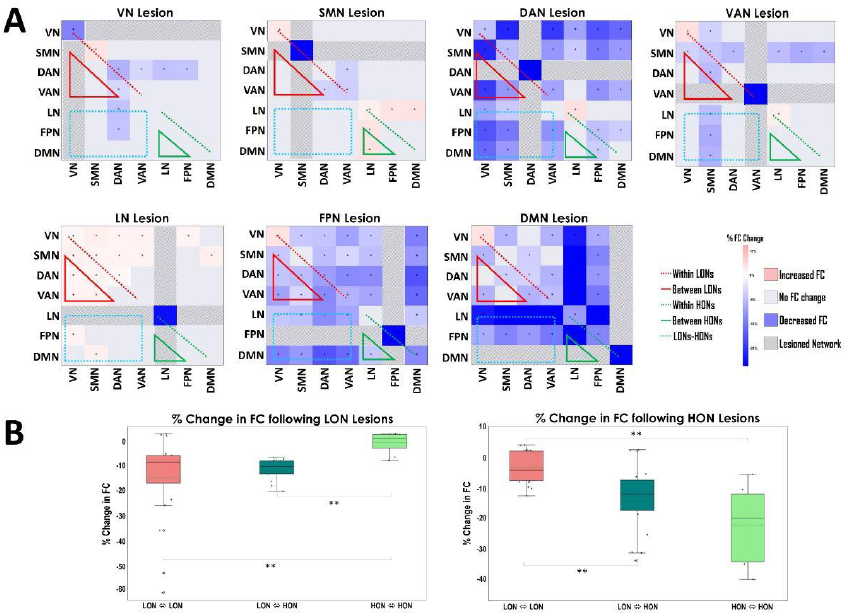
The effect of SC lesions on the FC of canonical networks for the WM Task. The legend for this figure is identical to Figure 4 for parts A and B. In this figure we can see the mutual antagonism between LONs and HONs is partially reversed in the WM task paradigm. This is particularly evident across the DAN, FPN, and DMN.

**Figure 7:**
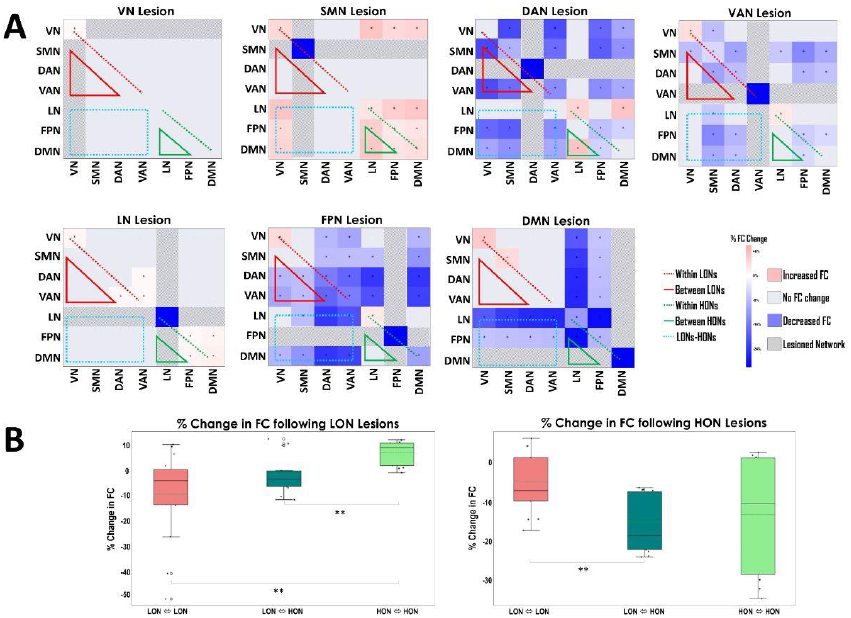
The effect of SC lesions on the FC of canonical networks for the Language Task. The legend for this figure is identical to Figure 4 for parts A and B. In this figure we can see the mutual antagonism between LONs and HONs is partially reversed in the language task paradigm. This is particularly evident across the DAN, VAN, and FPN.

The VN has widespread effects when lesioned in the resting state condition, but the effects of lesioning the VN are not as potent in task conditions. Furthermore, in task conditions, VN lesions tend to increase the activity of some LONs. In task conditions, SMN lesions tend to increase rather than decrease the FC of the VN and the FC between the VN and other networks. DAN lesions have the most robust effects out of all LON lesions and consistently decrease the FC within other LONs, and between LONs and HONs in all conditions. However, in WM and language task conditions, lesioning the DAN can also reduce the FC in some HONs (FPN, DMN) while increasing the FC in others. With VAN lesions, we observed a unique relationship between the VAN and VN wherein the FC within the VN and between VN and other networks increases following a VAN lesion in all conditions. The unique effect of each LON network lesion on different networks is shown in Figures 4 (resting state), 5 (motor), 6 (WM), and 7 (language) in panel A (***top***).

### HON SC lesions tend to have network and task-specific effects on the FC in LONs and HONs

Concerning HON lesions, there is a general decrease in the FC within and between HONs (15.3% average reduction). Like the lesioned LON, the lesioned HON typically has the most severe decrease in FC (sometimes above 30%). Additionally, there is a decrease in the FC between LONs and HONs (13.9% average reduction). Interestingly, following HON lesions, there are two generic groups of observations. The first includes the resting state (Fig. 4B [right]) and motor task (Fig. 5B [right]), where following any HON lesions, there is an overall FC increase within and between LONs (2.7% average increase). But for the other group, which involves the WM (Fig. 6B [right]) and language tasks (Fig. 7B [right]), following HON lesions, there is a decrease in the overall FC within and between LONs (4.6% average reduction). Again, the above results are a generalization. HON lesions also produce specific effects for each network.

When the FPN is lesioned, there are widespread effects on the FC of LONs and HONs. Consistent decreases in FC are seen in the DMN and LN and their respective connections to the other networks. In the resting state, FPN lesions increase the activity of some LONs, however, there is a widespread reduction in LON activity, following an FPN lesion, in the WM and language tasks. Following a DMN lesion, we observed a broad range of effects on the FC of LONs and HONs. Like the FPN lesions, DMN lesions tend to reduce the FCs of the FPN and LN and their connections to the other networks. However, except for the WM task, DMN lesions increase the activity of some or all LONs. In the WM task, DMN lesions reduce LON activity. There is an opposite effect on HON FC after an LN lesion. Unlike FPN and DMN lesions which reduce the FC of other HONs, LN lesions increase the activity of the DMN and FPN. Across all four conditions, there is a consistent increase in LON FC following an LN lesion in some or all LONs. The unique effect of each HON network lesion on different networks is shown in Figures 4 (resting state), 5 (motor), 6 (WM), and 7 (language) in panel A (***bottom***).

## Discussion

This study used a whole brain modelling approach to study the relationship between SC and FC in resting state and task-based fMRI conditions. Our results offer novel insights into the link between functional brain dynamics and the underlying anatomical connectivity as a function of a given brain state (i.e., resting state or task). In broader terms, these findings serve as a proof-of-concept, demonstrating the feasibility of studying the effects of SC lesions on FC across different conditions using a comprehensive whole-brain modelling approach. While these observations are intriguing, we must acknowledge several significant caveats and nuances in their interpretation, which we discuss below.

### SC network lesions induce local and global network FC changes

Network SC lesions influence the FC within the lesioned network and its relationships with other networks in the brain. In every condition (resting state or task), lesioning a LON or HON leads to a significant decrease in the FC of that network. However, the effect on the other networks varies depending on the type of network, and the condition being studied. While there are exceptions, typically, LON lesions reduce the FC within and between other LONs, and HON lesions decrease the average FC within and between all HONs, regardless of the condition.

Previous studies have used a similar approach to study the effect of lesions on FC. For example, one study observed the effects of lesions in the SC matrix in the macaque cortex (Honey & Sporns, 2008). The authors report that lesions involving high-degree nodes (i.e., nodes with a higher number of edges) had the largest effect on simulated cortico-cortical FC. In another study (Alstott et al., 2009), these concepts were extended to simulate BOLD FC using a conductance-based neural mass model. This study used diffusion spectrum imaging to obtain the SC matrix that included 66 anatomical regions subdivided into 998 parcels. The study reports that the FC is resistant to random lesions but highly susceptible to lesions that target nodes based on their centrality. Node centrality refers to the influence a node has within a network. Moreover, the authors observed non-local effects of the lesions, indicating changes in FC in ROIs distant from the lesion site. Subsequent analysis revealed that the extent of lost connections due to the lesion served as a reliable predictor of global FC changes. A different study (Váša et al., 2015) used the Kuramoto model to simulate fMRI data, and graph theory metrics to evaluate the effects of node lesions on FC and metastability. The removal of ‘hub’ nodes and nodes with high centrality led to a decrease in global FC, with global metastability increasing as a result of the lesions. Locally, there was a decrease in the FC following the removal of nodes with the node centrality and strength primarily determining the negative change in FC. Recently, an experimental study investigating DMN network organization in the rat brain utilized reversible chemical lesions in the dorsal anterior cingulate cortex (dACC) (Tu et al., 2021). These lesions significantly decreased DMN functional connectivity by reducing connectivity between network nodes. Consequently, changes in DMN connectivity induced by the lesion correlated with observable behavioral changes in the animal. This approach underscores the potential utility of targeted node lesions in generating preclinical models for studying brain disorders associated with network dysfunction.

The results from the present work draw parallels from those of previous studies. Instead of removing individual nodes, we structurally isolated entire networks by removing their connections to all other networks. This way, the relationship between networks based on the brain state is studied as a function of their underlying SC. We extend the previous work by investigating how these networks interact with each other when one of them is lesioned, and importantly, how this relationship changes based on a given condition (i.e., resting-state or task). For example, a lesion of VN does not have the same effect in resting state compared to task conditions. When the VN is lesioned in the resting state, other LONs see a decrease in average FC, but HONs see an increase in their average FC. However, in tasks, the effects of VN lesions are not as potent on LONs or HONs. LONs and HONs differ in both their functions and anatomical locations. By looking at how the FC interaction between networks changes based on the brain state helps us understand how brain networks utilize SC during certain conditions but not others. Based on our results, brain states that necessitate the integration of neural processes across multiple functional networks appear to rely more heavily on SC (Fukushima et al., 2018)

### LONs and HONs display a mutual antagonism in the resting state

We observed a mutually antagonistic relationship between LONs and HONs in the resting state. In other words, generally, when any LON is lesioned, there is an increase in the activity within and between HONs, and the converse also holds. For example, a DAN lesion leads to an increase in activity within and between the DMN, FPN, and LN. In contrast, when the DMN is lesioned, there is an increase in the average FC within and between the DAN, VAN, SMN, and VN (Fig. 4A).

A significant amount of research has studied network organization in the resting state, the majority of which has focused on the DMN (Mancuso et al., 2022). The earliest reports of the DMN describe it as a network specifically associated with the resting state. These so-called ‘task-induced deactivations’ highlighted areas of the brain that were consistently inactive in task conditions and were active at rest (Shulman et al., 1997). The brain’s metabolism was uniform when no specific task was being performed. Thus, this state of rest was considered to be the brain’s default activity level and the brain regions associated with it came to be called the DMN (Greicius et al., 2003; Raichle et al., 2001; Raichle & Snyder, 2007). However, as these concepts evolved, it became clear that the ‘resting’ brain is not in fact at rest. Rather, the brain is involved in activities such as imagining the future, recalling the past, thinking about others, and imaginary situations, collectively referred to as ‘mind wandering’ (K. C. R. Fox et al., 2015). Subsequent studies have shown that the DMN is typically anticorrelated with the DAN (M. D. Fox et al., 2005; Raichle, 2015; Uddin et al., 2009). There is evidence to support the notion that this anticorrelation between the DMN and DAN is driven, at least partially, by inhibitory interactions (Greicius et al., 2003; Knight et al., 1999). Additionally, research suggests that such anticorrelations are caused by ensemble inhibition, which refers to the coordinated activity of inhibitory neurons regulating the interactions between brain areas (Nyberg et al., 1996). If so, a plausible explanation for our results is that SC lesions of either network (DMN or DAN) remove this inhibition and the resulting disinhibition leads to an increase in FC of the other network.

Furthermore, studies have shown that the DMN and FPN are positively correlated at rest. There is evidence that the DMN and FPN embody overarching neural systems within the brain, composed of multiple sub-networks that engage in interactive dynamics (Marek & Dosenbach, 2018). This is consistent with what we see in lesions, where a lesion of the FPN also leads to an increase in the FC of some LONs and vice versa.

### LONs and HONs display more coordination during WM and language task conditions than rest

In the WM (Fig. 6A) and language (Fig. 7A) task conditions, the above relationship between LONs and HONs changes. While lesioning LONs still leads to an increase in the average FC in some HONs, we observe a reduction in some specific HON interactions. Moreover, lesioning HONs in task conditions (especially the FPN) leads to a widespread decrease within and between LON FC. Thus, in task conditions, the mutual antagonism between LONs and HONs at rest is no longer observed.

More recently, it has been shown that the DMN is not exclusively a ‘resting state’ network. Many studies have shown contrary results to the aforementioned ‘task-rest’ dichotomy of the DMN (Mancuso et al., 2022). For example, studies have shown that the DMN is responsible for stimulus-driven thoughts in task-based paradigms to improve task outcomes (Sormaz et al., 2018). The DMN’s implication in creativity (Huo et al., 2020) and problem-solving (Gerlach et al., 2011) also suggests its involvement in enhancing cognitive control during tasks necessitating internal thought processes. It has been proposed that the DMN likely fulfills these task-related functions in conjunction with the FPN (Gerlach et al., 2014; Spreng et al., 2014). In addition to activation studies, FC of the DMN with the FPN, and the attention networks increased in task paradigms like WM (Koshino et al., 2011, 2014; Murphy et al., 2020) and semantic memory retrieval (Fornito et al., 2012). Thus, substantial evidence indicates the essential role of DMN functionality, not only in internal mind-wandering activities mentioned above but also in the performance of external activities like WM.

The FPN is considered a functional hub for cognitive control due to its strong connectivity with other brain networks and likely plays a crucial role in task adaptation and implementation. For example, multiple overlapping FPN circuits are involved in WM (Murray et al., 2017) in addition to the FPN-DMN coordination mentioned earlier. Additionally, research shows that the FPN is critical for context-dependent language comprehension (Smirnov et al., 2014) and processing (Tomasi & Volkow, 2020). The strength of functional integration of the FPN with the rest of the brain positively correlates with overall cognitive ability, highlighting its significance in supporting superior cognitive functioning. These findings suggest that the FPN is involved in initiating and flexibly modulating cognitive control, enhancing the human brain’s ability to adapt (Marek & Dosenbach, 2018).

There are two primary attention networks, namely the DAN and VAN (Corbetta & Shulman, 2002). The DAN is typically associated with top-down attention concerning goal-directed tasks. This allows the DAN to use the task instructions or goal requirements to control task execution or carry out goal-directed behaviors (Vossel et al., 2014). The DAN has been consistently implicated in WM tasks across multiple modalities (Majerus et al., 2018). Importantly, DAN activity is directly proportional to the load of WM conditions, with higher WM loads leading to maximal DAN activity. For language, the DAN is also implicated in speech production. Research has shown that the DAN is responsible for a motor-sound interface that maps motor representations of speech sounds for articulation (Cloutman, 2013). On the other hand, the VAN is concerned with bottom-up attention that is driven primarily by external stimuli. This allows the VAN to support the detection of unexpected or unattended stimuli, allowing for a shift of attention (Vossel et al., 2014). The VAN is involved in the understanding of both spoken and written language by converting sound and/or word to meaning. Taken together, the DAN and VAN are important networks for language production and comprehension (Cloutman, 2013).

Our results show that in the WM task, DAN lesions reduce the average FC in the DMN and FPN. Lesions in the DMN or FPN lead to a decrease in the average FC across multiple LONs, including the DAN. All three networks (DAN, DMN, and FPN) are known to be involved in WM in either the planning (DMN) or execution (DAN, FPN). Therefore, lesioning one of these networks leads to a corresponding decrease in the others as seen in our results.

For the language task, we have seen how the DAN, VAN, and FPN are critical for language production, understanding, and processing, respectively. Our results show that lesioning either one of these three networks leads to a decrease in the average FC within the other two, as well as between the other two and the other functional networks not directly involved in language. In this way, the mutual antagonism between the DAN/VAN (LONs) and FPN/DMN (HONs) in the resting state is reversed in the WM and language task conditions.

### Cognitive load impacts network relationships following SC lesions

While the WM and language tasks show a reversal in LON-HON relationship, it is interesting to note that tasks with a ‘lower cognitive load’ do not see this reversal of antagonism. We refer to the motor task which involves basic movement of the fingers, toes, or tongue based on a visual cue (see *Methods* above). The SMN was one of the earliest resting-state networks to be described (Biswal et al., 1995) and is involved in the voluntary motor and somatosensory processes concerning the limbs, body, head, neck, and face (Seitzman et al., 2019). Previous research has shown that increasing the complexity of motor tasks leads to the recruitment of other brain regions from multiple different networks (Bullmore et al., 2003; Hummel et al., 2003; Puh et al., 2007). Seeing as how the motor task data used for this study was acquired from a relatively simple set of tasks/instructions, an argument can be made that this task exclusively engages the SMN without involving the other networks (Gabitov et al., 2019; Shine et al., 2016). Therefore, tasks with a lower cognitive load like the motor tasks potentially share a similar network connectivity profile to the resting state (which by definition has a low or no cognitive load). When compared to WM or language tasks which have a higher cognitive load, we do not see a reversal of antagonism between LONs and HONs following SC lesions in motor tasks. Lesioning LONs (or HONs) in the motor task leads to an increase in the average FC of HONs (or LONs). In this regard, motor task SC lesions have similar effects to what we see with SC lesions in the resting state (described above).

### Limitations

Although the findings of this study offer valuable insights into the interactions among brain networks following SC network lesions in both resting-state and task-based conditions, it is important to note some significant limitations.

#### Simplified Models

In our study, we employed the RWW model, which is a simplified mathematical representation of brain dynamics. While this model serves as a useful tool for exploring certain aspects of brain function, it does not encompass the full complexity and diversity observed in real brain networks. Real brain networks exhibit various features, including oscillations, chaotic behavior, and the generation of spatiotemporal patterns, which are not fully captured by the RWW model. Consequently, the simplifications and assumptions inherent in the model may restrict the applicability of our findings to real-world brain network interactions.

#### Model Fit to Empirical Data

The average fit of our model-based simulations to their empirical counterparts is 0.64 for the resting state condition, and 0.53, 0.49, and 0.31 for the motor, WM, and language tasks, respectively, across our 200 subject cohort. These values fall around previously established model fits to empirical data, which range from 0.1 to 0.5 (Messé et al., 2015; Roberts et al., 2019). We speculate that this difference in the fit between resting-state and task data is primarily due to the length of the empirical fMRI time series associated with each condition. As per the HCP database, the resting-state time series comprises 1200 time points, the motor task time series comprises 284 time points, the WM time series comprises 405 time points, and the language time series comprises 316 time points.

Given that the resting-state time series contains roughly three to four times as many time points as the task conditions, it provides a richer temporal context for model training and parameter tuning using the RWW model. With a longer time series, the ANN has more opportunities to capture subtle temporal dynamics and patterns inherent in the data. This increased sample size enables the ANN to learn the underlying structure of the neural signals and refine its model parameters more effectively. Consequently, the longer time series facilitates a more comprehensive exploration of the complex temporal dynamics represented by the RWW model, enhancing the model’s ability to accurately represent and predict brain activity patterns.

#### Node Lesions instead of Network Lesions

Previous works (Alstott et al., 2009; Honey & Sporns, 2008; Tu et al., 2021) have studied the effects of lesioning specific nodes within brain networks to study their effects on network FC and how it changes as a result of the lesion (For example, lesioning an important node/hub within the DMN). In this study, we structurally isolated the entire network to observe its interactions in resting-state versus task-based conditions. However, this approach did not allow us to study the functional architecture within the network or how it changes in response to structural isolation. Consequently, we could not examine interactions between different regions within a network at the node level. Combining the methodologies presented in this study with previous works, subsequent research can study the effects of lesioning multiple nodes within a given network and study the changes in network behavior in the different brain states.

#### Potential Clinical Applications

All of the fMRI data we used in this study were obtained from 200 healthy participants from the HCP database. Network and node SC lesions can be used to mimic pathological conditions such as tumors or strokes (Váša et al., 2015). In this case, the pathology dictates the extent of the lesion, and analysis can be carried out in a similar fashion presented above. This would allow us to better understand the network interactions in such pathophysiological conditions. For example, previous work has strongly suggested that disrupting SC locally or globally leads to alterations in FC that resemble the FC of schizophrenia patients at rest (Cabral et al., 2012). Thus, there is a need for further investigation into the unique SC-FC interplay and its corresponding changes in neuropathological conditions.

## Conclusions

This study employed the RWW whole brain model within a PyTorch framework to explore the intricate interplay between SC and FC in resting state and task-based fMRI conditions. Our findings shed light on how functional brain dynamics are intertwined with underlying anatomical connectivity, showcasing the adaptability of these relationships across different brain states. We observed that SC network lesions induce significant changes in overall network connectivity, impacting FC within the lesioned network as well as its interactions with other brain networks. Lesions affecting LONs and HONs exhibit distinct effects on FC, with LON lesions generally diminishing FC within and between other LONs, and vice versa for HON lesions, albeit with some exceptions contingent upon network type and condition. Notably, mutual antagonism between LONs and HONs is evident in the resting state, where lesioning one network type results in heightened activity within and between the other network type, a relationship that reverses in task conditions. In the WM and language tasks, this antagonism diminishes, leading to more coordinated interactions between LONs and HONs. However, tasks with lower cognitive demand, such as motor tasks, display network connectivity profiles akin to the resting state, exhibiting no reversal of antagonism following SC lesions. Our findings underscore the dynamic nature of brain network interactions, highlighting their modulation by different brain states and task demands. Additionally, these insights hold promise for potential clinical applications, offering valuable implications for understanding and managing conditions such as brain tumors and stroke, where the pathology leads to an SC lesion, ultimately informing personalized treatment strategies and rehabilitation approaches.

## Supplementary Information

### Permutation Analysis

We start with a 7×7 matrix, where each element represents the observed changes in the average FC of the given network to the other networks (represented as % changes; see Figs. 4-7) when one of the seven networks is lesioned, in resting-state and task fMRI simulations. As these matrices are symmetric, the upper triangle values are set to 0. These values represent the observed test statistic and will be used for comparison against shuffled data. The non-zero values from the diagonal and lower triangle of the matrix are permuted randomly. This permutation process is repeated for a large number of iterations (10,000 times), generating a distribution of shuffled values under the null hypothesis. In our study, we applied this permutation testing procedure to examine the significance of patterns observed in 7×7 matrices. By comparing the original results to the distribution generated from shuffled permutations, we assessed whether the observed percent changes in 7×7 matrices were statistically significant or could be attributed to random variation.

**Supplementary Figure 1:**
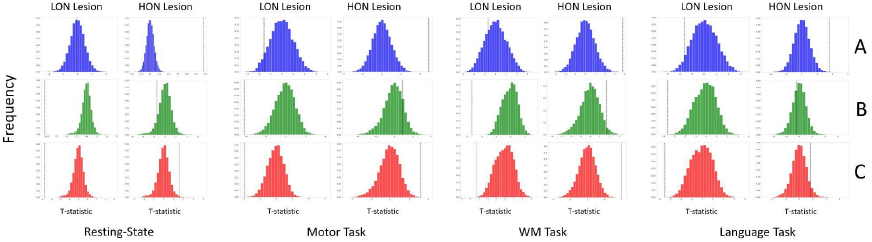
Permutation analysis of FC changes. To verify the results obtained in Figs. 4-7 – B, we carried out a permutation analysis for all four conditions. A: histogram of t-statistic between LONs and LON-HON; B: histogram of t-statistic between LON-HON and HONs; C: histogram of t-statistic between LONs and HONs. The left and right column for each condition represents LON and HON lesions, respectively. The histogram plots show that the distribution of t-statistic values following the randomized shuffled permutations of the FC values in the 7×7 grids described earlier. The originally observed t-statistic is marked witht black dotted vertical line. The significance of the original t-statistic is determined by whether or not it falls within the histogram of the t-statistics following the randomized shuffled permutations. If the black dotted veritcal line falls within the histogram, the original t-statistic is not significant. If the black dotted vertical line is outside or at the edge of the histogram, then it is significant at either p < 0.01 or p < 0.05, respectively. Refer to Figs. 4-7 B for significance values.

### RWW, Balloon model, and Fitted value tables

Additional information regarding default and estimated model parameter values can be found below.

